# Global shark species richness is more constrained by energy than evolutionary history

**DOI:** 10.1101/2022.04.15.488537

**Authors:** Emmaline R. Sheahan, Gavin J.P. Naylor, Daniel J. McGlinn

**Affiliations:** Department of Biology, College of Charleston, Charleston SC; University of Florida

**Keywords:** biodiversity, biogeography, diversification, energy gradient, temperature gradient, niche conservatism, simulation

## Abstract

**Aim:** To examine the support of two ecological diversity theories- The Ecological Limits Hypothesis (ELH) and the Niche Conservatism Hypothesis (NCH) - in explaining patterns of global shark diversity.

**Location:** Global scale and two ecological realms: the Tropical Atlantic and the Central Indo-Pacific.

**Time Period:** Past 100 years

**Major Taxa Studied:** We examined 534 species of sharks and chimaeras, and we performed two subclade analyses on 272 species of ground sharks and 15 species of mackerel sharks.

**Methods:** We compared the species richness, mean root distance (MRD), and tree imbalance patterns to those simulated under the ELH and NCH with temperate and tropical centers of origin. We used sea temperature as a proxy for energy availability. We examined the importance of biogeographic history by comparing the model fits between two taxonomic groups, ground and mackerel sharks, and two geographic regions, the Tropical Atlantic realm and Central Indo-Pacific realm.

**Results:** The ELH, temperate-origin model had the best fit to the global dataset and the sub-analyses on ground sharks, mackerel sharks, and the Tropical Atlantic. The NCH temperate-origin model provided the best fit for the Central Indo-Pacific. The β metric of tree symmetry showed the best potential for differentiating between the ELH and NCH models, and the correlation coefficient for temperature vs MRD performed the best at differentiating between temperate and tropical origin of ancestors.

**Main Conclusions:** The global and subclade analyses indicate the ELH provides the best explanation for global scale shark diversity gradients even in clades with varying ecology. However, at the realm scale, biogeographic history has an impact on richness patterns. Comparing multiple metrics in relation to a simulation model provides a more rigorous comparison of these models than simple regression fits.

## Introduction

Biodiversity gradients have captured scientists’ imaginations for over 150 years, yet we still are not much closer to clearly accepted explanations for some of the largest and most striking patterns, such as the latitudinal diversity gradient. The latitudinal diversity gradient is a well-documented phenomenon that holds across a myriad of terrestrial and marine taxa in which richness appears to be highest at lower, warmer latitudes and declines as latitude increases towards the poles (Rosenzweig., 1995). One of the difficulties with studying this phenomenon is that there are many hypotheses to explain diversity gradients, but they often involve similar mechanisms or assert very similar predictions, making it difficult to differentiate their support empirically (Palmer, 1994, Hulbert and Stegen 2014a,b, Pontarp et al. 2018). Many of these hypotheses relate directly to the mechanisms which control species accumulation in an area-such as speciation, extinction, and dispersal. The speciation hypothesis asserts that speciation and extinction rates differ latitudinally (Willig, 2003), while the time-for-speciation hypothesis (Rabosky, 2011) holds that species originating in the tropics have had more time to accumulate there and dispersed into temperate regions later. The disturbance hypothesis holds that extinction is greater at higher latitudes due to an increased incidence of disturbance related to glaciation in temperate regions while the tropics have been relatively untouched by this effect (Fraser & Currie., 1996). Aside from these evolutionary and geographic mechanisms, others have posited that the latitudinal diversity gradient is the result of the amount of energy available to a system in the form of temperature or net primary productivity (Francis & Currie., 2003), and that these resource limitations subsequently lead either to dispersal, extinction, or sympatric speciation. It is helpful to simplify this range of hypotheses into two groups: the Ecological Limits Hypothesis (ELH) and the Niche Conservatism Hypothesis (NCH).

The NCH has three primary components (Wiens et al., 2006): (1) that richness will be highest in habitats nearest to the ancestral niche because range of tolerance is phylogenetically conserved, thus making dispersal into temperate zones difficult, (2) there is more time for species divergence in areas already inhabited relative to regions that depend on immigration, and (3) that tropical regions occupied a larger geographic area until relatively recently, thus allowing for more species (Rosenzweig, 1995; Hawkins & Porter, 2001). The ultimate consequence of this is a gradient of species richness where in the most diverse areas are those habitats which best reflect the ancestral niche, and diversity decreases the further one moves away from these areas (Wiens & Donoghue, 2004). The NCH is primarily concerned with evolutionary factors and allows for species to accumulate indefinitely. In contrast, the ELH holds that the total number of species that can inhabit an area is limited by available energy due to competition for limited resources. This idea is related to the concept of species carrying capacity and the notion that there has not been a clear trend in species diversity through geologic time, calling into question the assumption that species richness increases indefinitely through time (Rabosky, 2009). Modeling work done by Etienne et al (2019) suggested that energy constraints play a large role in determining global patterns of richness, and that other competing forces are only influential before equilibrium is reached.

Although the ELH and NCH are based on different mechanisms, they produce similar richness gradients if regions of high energy coincide with regions identified as the ancestral climate niche for a clade (e.g., tropical biomes). If the relationship between species richness and a spatial gradient is the only factor considered, it is not possible to discern which hypothesis better explains the pattern. Hurlbert and Stegen (2014a,b, referred to as H&S hereafter) developed an eco-evolutionary simulation framework that provides additional predictions beyond just a simple increase in richness with energy and thus a stronger test for distinguishing between the ELH and NCH. Specifically, H&S ran simulations where species evolved along an energy gradient through time following either the NCH or ELH. The ELH simulations enforced a species carrying capacity that limited how many species could coexist at a given energy level. H&S also varied whether the ancestral niche was in the temperate (low energy) or tropical (high energy) region. H&S used several metrics to differentiate between these four potential biogeographic pathways: species richness, phylogenetic tree symmetry (β) (Blum *et al*., 2006), and a measure of how phylogenetically derived co-occurring species are using mean root distance (MRD) which is the average number of nodes from root to tip (Kerr & Currie., 1999). The H&S simulation can be used to uncover the following predictions (Fig. 1, summarized in Table 1):

**Figure 1.**
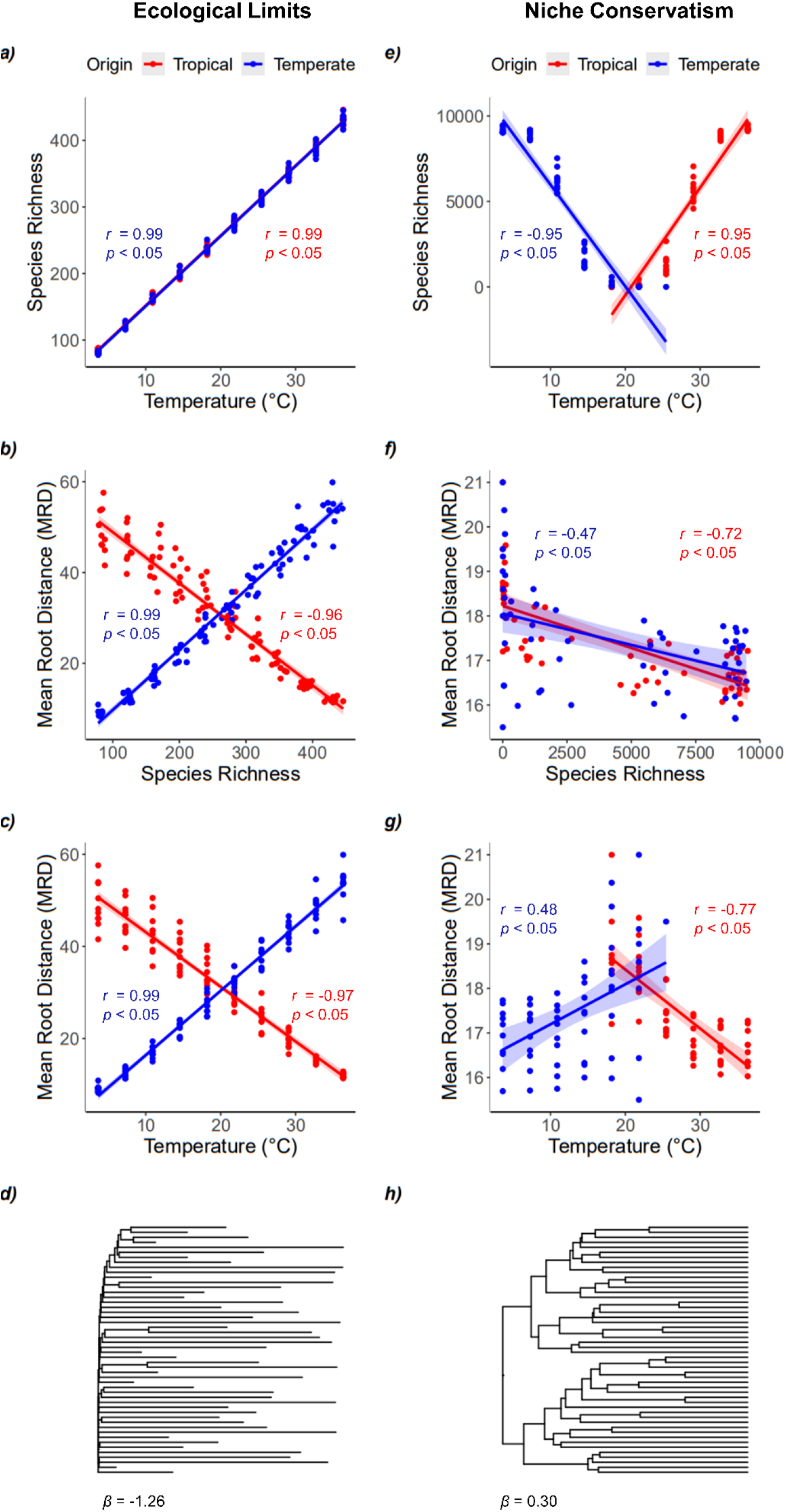
The results of the H&S simulation model in which temperature is used as a proxy for energy for the Energy Limits Hypothesis (a - d) and for the Niche Conservatism Hypothesis (e - h). Results with a tropical center of origin are shown in red and results from a temperate center of origin are shown in blue. Best fit lines are simple linear regression lines with 95% confidence intervals. We also report Pearson correlation coefficients (*r*) and associated *p*-values for each linear fit. The phylogenetic trees displayed in this figure were randomly pruned to 50 tips for visual clarity, but the estimated tree symmetry (β) used all tips. Because the climatic origin did not influence the symmetry of the phylogenetic trees (as can be observed in figure 3), only one representative tree is shown for ELH (g) and NCH (h) respectively.

**Table 1.** The expected directionality of the relationships between temperature- used in our analysis as a proxy for energy-, richness, and MRD, and whether the phylogenetic tree is balanced (+) or unbalanced (-) for the NCH and ELH of either a tropical or temperate origin generated using the H&S model.

| Model | Temperature<br>vs Richness | Richness vs<br>MRD | Temperature<br>vs MRD | Tree Symmetry<br>($\beta$) |
| --- | --- | --- | --- | --- |
| ELH Temperate | + | + | + | — |
| ELH Tropical | + | — | — | — |
| NCH Temperate | — | — | + | + |
| NCH Tropical | + | — | — | + |

The ELH predicts:

1. The correlation between energy and richness is positive for both centers of origin (Fig. 1a).
2. The correlation between phylogenetic derivation (i.e., MRD) and species richness is negative for a tropical origin and positive for a temperate origin (Fig. 1b).
3. The correlation between energy and MRD will be negative for a tropical origin and positive for a temperate origin (Fig. 1c).
4. Less balanced phylogenetic trees than randomly expected (β < 0) for both centers of origin (Fig. 1d).

The NCH predicts:

1. The correlation between energy and richness is positive for a tropical origin and negative for a temperate origin (Fig. 1e).
2. The correlation between MRD and regional species richness is negative for both centers of origin (Fig. 1f).
3. The correlation between energy and MRD will be negative for a tropical origin and positive for a temperate origin (Fig. 1g).
4. More balanced phylogenetic trees than expected (β > 0) for both centers of origin (Fig. 1h).

The simulation framework created by H&S, though still quite qualitative and overly simplified, is an attempt at formulating stronger methods of testing the ELH and NCH by examining multiple relationships simultaneously (Pontarp et al 2018). In a similar vein, this study aims to bridge the gap between qualitative and quantitative methods by applying the H&S framework to understand the global richness gradients for sharks.

Sharks represent an interesting test case for the ELH and NCH because these hypotheses are often compared using species that diversified during the tropical portion of the Eocene (e.g., mammals, birds, flowering plants, Wiens and Donoghue 2004). Sharks, by contrast, are an old clade extending back 450 million years (Grogan & Lund, 2012). The majority of modern shark diversification occurred from the Jurassic to the early Cretaceous (Sorenson *et al*., 2014). Though the Earth was considered to be in a greenhouse period during this time with relatively low climate gradation and warm ocean temperatures extending into high latitudes (Jeynkins *et al*., 2012), this was a part of a warming trend which peaked with the Paleogene-Eocene Thermal Maximum. Additionally, modern ocean basins and continental arrangements had not yet fully formed (Brunetti *et al*., 2015). The range in environmental variation both ancient and modern among chondrichthyans in conjunction with their evolutionary persistence lends the group to an assessment of the forces that have shaped their diversity over time. Sharks are particularly interesting because their biogeography does not follow the patterns widely ascribed to groups whose spatial distributions are consistent with the NCH. By contrast, shark richness appears to be largely coastal, with a few specific hotspots where their diversity is especially high, such as the coasts of South Africa, Eastern and Western Australia, and Southern Japan (supplemental figure 1). Additionally, sharks are ecologically important as top-down regulators in most marine habitats and are consequently of high conservation concern. Understanding the mechanisms that govern shark biodiversity at the global scale can guide policy directed toward preserving this ancient and significant clade.

The purpose of this study is to estimate the support of the ELH and NCH for explaining global shark diversity. Additionally, we examine the generality of that support by examining two biogeographic realms and two ecologically divergent subclades.

## Methods

### Simulation analysis

We used the eco-evolutionary simulation model of H&S to derive the four expected biodiversity patterns described in the Introduction under ELH and NCH with temperate and tropical centers of origin. We used the default parameters, wherein under the NCH temperature along the gradient varied from 0 – 40 °C, the regional carrying capacity throughout was 40,000 individuals, and the per individual speciation probability was 10^-6^. Under the ELH every parameter was the same except for the regional carrying capacity, which varied along the gradient from 4,000 – 40,000 individuals. The R code to run the H&S simulation is available here: https://github.com/ahhurlbert/species-energy-simulation

We modified code from the function `regional. calc` in the `reg_calc_and_analysis.R` script defined at the above repository to calculate MRD (https://doi.org/10.5281/zenodo.5523198), and we computed β using `apTreeshape::maxlik.betasplit` (Bortolussi, 2012) with a profile confidence interval. We then generated linear plots for temperature versus species richness, species richness versus MRD, and MRD versus temperature, as well as output the associated Pearson correlation coefficients, p-values, and confidence intervals.

### Empirical analysis

We mapped and summed range data of 534 shark species accrued and updated by Naylor et al (2012) and published by the IUCN across a global gradient in the cylindrical equal area projection; this is a separate data set from that which has been garnered by the Census for Marine Life, a source which was used in an analysis by Tittensor et al (2010). The two datasets are likely similar, but the data set retrieved from Naylor and the IUCN is more recent and, we believe, likely more accurate due to the fact that species identifications were corroborated with genetic evidence. All original range maps can be accessed via Zenodo (https://doi.org/10.5281/zenodo.6321610). We rasterized the range data at six different spatial grains: 110 x 110 km, 220 x 220 km, 440 x 440 km, 880 x 880 km, 1760 x 1760 km, and 3520 x 3520 km. We did not find strong evidence of scale-dependence in the results of the analysis; therefore, we only report the results at the 880 x 880 km grain. Additionally, Tittensor et al (2010) examined this same spatial grain for species richness of various marine clades. We examined several proxy variables of energy for sharks: ocean chlorophyll (NASA Earth Observations, https://neo.sci.gsfc.nasa.gov/view.php?datasetId=MY1DMM_CHLORA), temperature, and bathymetry (NOAA’s World Ocean Database, https://www.nodc.noaa.gov/OC5/WOD/pr_wod.html). We ultimately used temperature as our environmental energy gradient given its strong correlation to shark richness relative to bathymetry and chlorophyll (Supplemental figure 4). Due to the broad range of depths inhabited by sharks, we used temperature based upon the mean from the surface down to 3700 meters, the lowest depth at which any shark species is known to occur (Priede *et al*., 2006). The analyses also included a phylogenetic tree compiled by Naylor et al (2012). The tree excluded 275 species due to inadequate data, and we resolved polytomies by averaging the branch lengths of each involved tip.

With the empirical global range map data, we calculated species richness and MRD for all 534 shark and chimera species, and β for the 259 remaining species with adequate phylogenetic information. We also ran sub-analysis just for the 272 species of ground sharks (Carcharhiniformes) and the 15 species of mackerel sharks (Lamniformes). We compared these two clades because several species of mackerel shark have developed homeothermic regulation, an adaptation which has better allowed them to disperse into temperate and pelagic waters compared to the poikilothermic ground sharks (Legendre & Davesne, 2020). We were interested to see whether or not this trait difference would result in different outcomes between them. Finally, we carried out regional analyses on two different biological realms defined by WWF: the Tropical Atlantic and the Central Indo-Pacific (Spalding *et al*., 2007). We selected these two regions based on their geographic distinctness; while the Tropical Atlantic is comprised primarily of continental coastline, the Central Indo-Pacific contains many islands and is often referred to as a “species pump” for richness (Crame, 2001).

We extracted raw expected values for each metric for four different models-the ELH for species of a temperate origin, the ELH for species of a tropical origin, the NCH for species of a temperate origin, and the NCH for species of a tropical origin-from the H&S simulation. We procured the correlation coefficients and their associated p-values by running the `cor.test` function with the Pearson method for each pairing of metrics across all ten simulation repetitions. We averaged the beta values produced by phylogenetic trees across the repetitions for each model, though we dropped two trees from the NCH Tropical origin repetitions and 3 trees from both the ELH Tropical origin and the ELH Temperate origin. We had to drop these trees because they were too large for the `maxlik.betasplit` function to properly calculate. After plotting the observed versus the expected values for each metric for the global, subclade, and regional analyses, we calculated the mean sum of squares error (MSE) of every model for each analysis in order to garner goodness of fit. The code for the empirical analysis and comparison between the simulations and empirical data are archived here: https://github.com/mcglinnlab/shark-ray-div.

## Results

Global shark species richness was strongly positively correlated with ocean temperature (Fig. 2a). Mean Root Distance (MRD) was positively correlated with both richness (Fig. 2b) and temperature (Fig. 2c). The residuals of the linear fits showed systematic deviation for all three relationships which suggests that their linear approximation is coarse despite relatively high correlation coefficients (all r > 0.45). The phylogenetic tree of global sharks was highly unbalanced (β = −0.87, Fig. 2d) which certain clades showing more diversification (or less extinction) than others.

**Figure 2.**
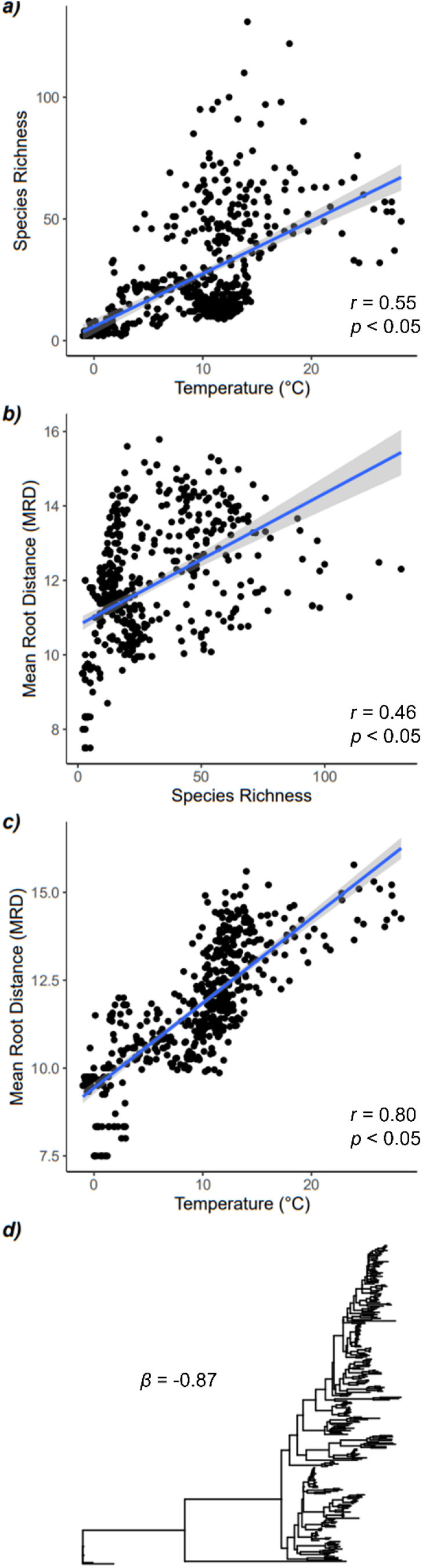
The empirically observed patterns of global sharks for: a) ocean temperature and species richness, b) species richness and MRD, c) ocean temperature and MRD, and d) phylogenetic tree symmetry. We also report Pearson correlation coefficients (*r*) and associated *p*-values for each linear fit. In panel d) the β statistic of tree symmetry is reported for the displayed phylogenetic tree.

The global empirical results were best fit by the Ecological Limits Hypothesis (ELH) for species of a temperate origin (Fig. 3, Table 2); however, the observed values for this scenario, with the exception of the β metric of tree imbalance, did fall consistently below the simulated predictions (blue points in Fig. 3a are all below the 1:1 line). The support for the ELH over the NCH depended in large part on the phylogenetic tree symmetry. The ELH predicted an unbalanced tree which is more congruent with what was observed empirically (Fig. 2d) while the NCH predicted a balanced tree. Within the context of the ELH, the support for the temperate origin was largely derived from the relationships between richness vs MRD and temperature vs MRD. The temperate origin ELH predicted these relationships would be positive as observed while the tropical origin ELH predicted negative relationships (Fig. 3a).

**Figure 3.**
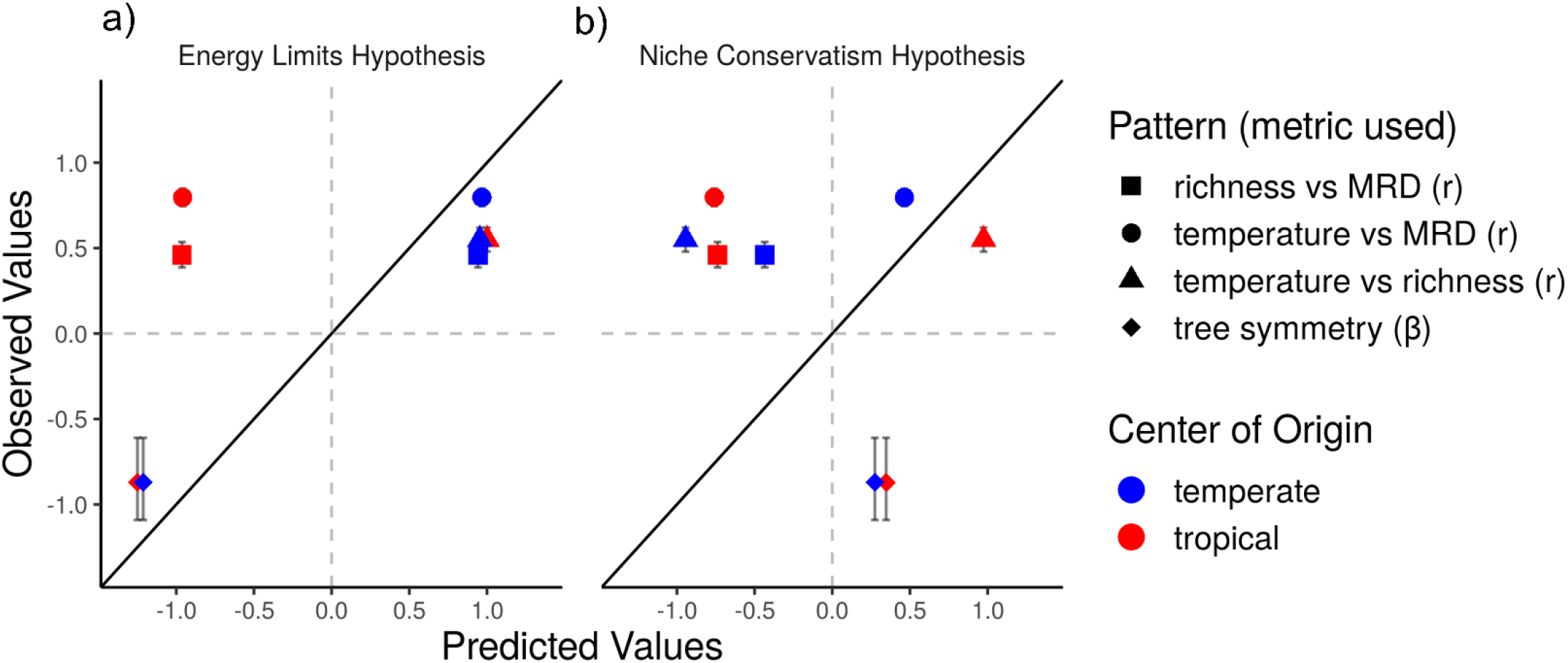
Predicted vs observed plots for the global analysis of the four metrics under the (a) Energy Limits Hypothesis (ELH) and (b) Niche Conservatism Hypothesis (NCH). The solid line is the one-to-one line; the closer the points are to the line, the better the fit between the model and reality. The 95% confidence interval of the observed metrics is provided for each metric, but it is sometimes obfuscated by the plotting symbol. The predicted values have been randomly jittered slightly to decrease point overlay. Points representing a tropical center of origin are red, and points representing a temperate center of origin are blue.

**Table 2.**
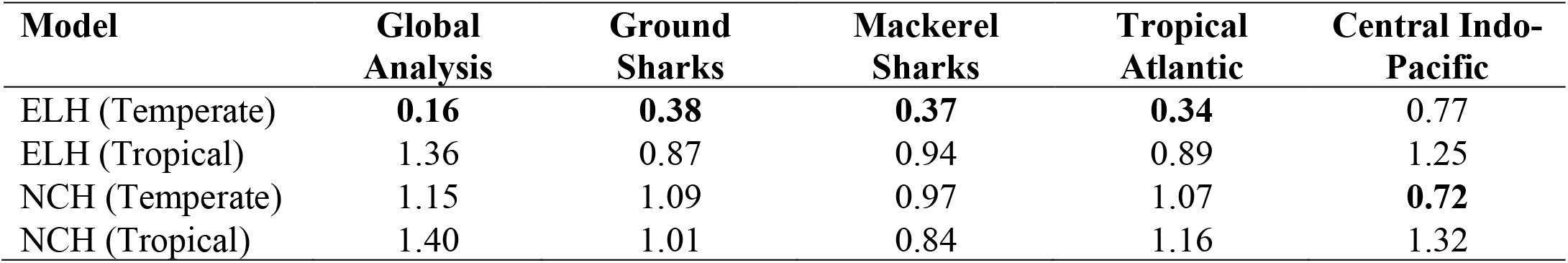
Mean sum of squares error (MSE) values between the global analyses and the sub-analyses and each simulated model. The lower the MSE, the better the model fits the observed pattern. The lowest MSE value for each analysis has been boldened. ELH of a temperate origin performs best for the global analysis, ground sharks, mackerel sharks, and the Tropical Atlantic. All MSE values are relatively high for the Central Indo-Pacific, with the NCH of a temperate origin performing the best.

The relationship between temperature and MRD appeared to have the most leverage on the support for the temperate vs tropical origin, and the metric of phylogenetic tree symmetry appeared most to support the ELH over the NCH (Fig. 3). Richness vs MRD (Fig. 2b, 0.46) exhibited a positive correlation as would be consistent with the ELH for species of temperate origin, and richness vs temperature (Fig. 2a, 0.55) was strongly positively correlated. While the relationship between MRD and species richness does well to differentiate the ELH of temperate origin from the other hypotheses, the correlation coefficient for MRD vs temperature exhibits greater disparities depending on region of origin itself (Fig. 3). The ELH for species of a temperate origin also exhibited a far lower mean sum of squares error (0.16) when applied to the global analysis than any other model (table 2). There is a large difference between the fit for the ELH for species of a temperate origin and the ELH for species of a tropical origin, far more so than between the associated versions of the NCH.

The ELH for species of a temperate origin also better represented both subclade analyses (table 2) than the other three models, a result again driven largely by the correlation coefficient for MRD vs temperature and the beta metric of tree imbalance. However, beta appears to be particularly sensitive to small trees, subsequently resulting in a far wider margin of error for the metric with regards to mackerel sharks, an order with only 15 species (supplemental figure 5). Consequently, the fit of any model for the mackerel shark order is less certain than that of the other analyses, all of which involve more than a hundred species and depict more stable beta values. While the observed vs predicted plot for the tropical Atlantic region appears consistent with both subclade analyses and the global analysis, the central Indo-Pacific region differs largely due to the negative relationship between species richness and temperature. The ELH for species of a temperate origin displayed the smallest mean squared error (table 2) for every sub analysis except for the Central Indo-Pacific, for which the NCH for species of a temperate origin displayed the lowest MSE (0.72). It should be noted however that all of the MSE values were relatively high and less varied for the Central Indo-Pacific, and the MSE for the ELH for species of a temperate origin (0.77) was not that much larger than that of the NCH for species of a temperate origin.

## Discussion

In the case of the global analysis, the ELH appears to have more of an influence in determining patterns of shark species richness than the NCH. Additionally, the ELH for species of a temperate origin appears to do better than the ELH for species of a tropical origin, a surprising result given the fact that it is believed that chondrichthyans originated in shallow tropical seas in the early Devonian period, and these early forms were typically small and fed at mid-trophic levels (Grogan & Lund., 2012). However, adaptive radiation in chondrichthyans during the Carboniferous period produced several larger bodied, apex feeding taxa capable of occupying a wider range of habitats, such as estuarine bays, epicontinental shelves, and deep and open ocean areas. Modern elasmobranchs are believed to be derived primarily from neoselachians, which included both coastal and trans-oceanic predators during the Jurassic and Cretaceous periods, and this clade is believed to be connected to Paleozoic forms via the hybodonts, which occupied a variety of marine environments (Grogan & Lund., 2012). The fact that modern elasmobranchs are primarily the result of clades which inhabited a variety of temperature and habitat regimes may account for the observed discrepancy between temperate and tropical origin.

Though H&S did not use the correlation between MRD and the energy gradient in their work, we included the metric due its ability to differentiate between models by climatic origin. Both the NCH and the ELH predict a positive correlation between MRD and the energy gradient for species of a temperate origin and a negative correlation for species of a tropical origin. These trends may possibly be explained by sympatric speciation. MRD is a measure of species derivation; the higher the MRD, the more highly derived species there are inhabiting an area. In terms of the ELH, the more energy there is in an area, the higher the organismal carrying capacity. For species of a tropical origin, because of the species carrying capacity, the tropics would fill up with less derived species, so there would be a higher proportion of more derived species as they moved into colder climate bands. For species of a temperate origin, the temperate region would fill up with less derived species, and there would be more niche space available for new species via speciation and niche partitioning in the tropics due to the higher carrying capacity. This results in a negative relationship between temperature and derivation for species of a tropical origin and a positive relationship between temperature and derivation for species of a temperate origin. This phenomena may also be explained in terms of metabolic rate: species inhabiting warmer climate regimes tend to express shorter generation times (McClain *et al*., 2012). Quian et al used the correlation between MRD and an environmental gradient as a proxy for diversification in a study aimed at explaining richness patterns in North American angiosperm trees (Quian *et al*., 2015), and while their results were negative regarding diversification rates driving richness, our observed data does display a positive correlation between MRD and temperature congruent with their expectation. Regarding the NCH, dispersal may be more difficult for lineages with narrow tolerances for environments outside of their conserved niche, resulting in an increase in MRD as temperate species move into tropical regions and as tropical species move into temperate regions. As a consequence, MRD would be positively correlated with temperature for species of a temperate origin and negatively correlated with temperature for species of a tropical origin. While something like a BAMM analysis on a time calibrated tree may be better able to establish the relationship between MRD and diversification rates, recent studies calling into question the validity of BAMM analyses due to the fact that extant time-trees are consistent with many equally possible diversification histories discouraged us from pursuing this avenue (Louca & Pennell., 2020).

The correlation between MRD and richness only differentiates between the ELH of a temperate origin and every other model. The simulation predicts a positive relationship between MRD and richness for species of a temperate origin and a negative relationship between species of a tropical origin for the ELH, while the NCH should, regardless of origin, yield a negative relationship. This can be explained for the NCH because richness is always highest in the region of origin, while MRD should be higher away from the region of origin because the species there are derived from older species which have dispersed out. This consequently creates an inverse relationship regardless of origin. In the case of the ELH for species of a tropical origin, richness is highest where there is more energy, in the tropics, but the energy constraint means that species can only accumulate there up to a certain limit, so all of the species that then begin to arise in the climatic bands moving to the temperate region are more derived. This results in a negative relationship between richness and MRD. For the ELH for species of a temperate origin, the temperate region would fill up with a smaller number of less derived species, and as species dispersed into the tropics, there would have been both more species due to the higher carrying capacity as well as more highly derived species. This would result in a positive relationship between richness and MRD. The energy constraint may also explain why the correlation is much stronger for the ELH compared to the NCH; without a species carrying capacity, more derived species can still appear in the region of origin even as other more derived species are appearing away from the region of origin due to dispersal, and this phenomenon is limited under the ELH due to the species carrying capacity. The ELH may produce negative beta values indicating unbalanced trees due to variable diversification rates, whereas the NCH is more closely related to dispersal and diversification rates subsequently remain consistent, producing relatively balanced trees with positive beta values.

The ELH for species of a temperate origin also better fit both subclade analyses than the other three models, though the large variation in beta for mackerel sharks makes this result somewhat more dubious than that of the global analysis. The mackerel and ground shark orders were specifically selected for comparison due to the novel adaptation in the former of homeothermic regulation, a property that has enabled this group to inhabit temperate and open seas, potentially reducing the influence of temperature on dispersal. However, this taxonomic difference does not seem to matter with regards to determining the influence of biogeographical hypotheses, seeing as both groups expressed the lowest MSE for the ELH for species of a temperate origin. The subregion analyses, however, do seem to suggest that regional geography has an impact in determining richness patterns. The tropical Atlantic closely resembles much of the area covered in the global analysis seeing as it is mostly comprised of continental coastline, and thus the fact that both best cohere with the same hypothesis could be expected. The Central Indo-Pacific, by contrast, is considered a species pump where historical vicariance events throughout the Cenozoic, such as tectonic activity and eustatic changes in sea level, are believed to have caused dramatic allopatric speciation (Crame., 2001). This is reflected by the fact that no model produced a very low MSE for the Central Indo-Pacific realm, subsequently indicating that biogeographical hypotheses applied at the global level may not be as important to determining patterns of richness at the regional level as geographic history.

## Conclusions

We conclude based upon the positive correlations observed between temperature and shark taxonomic richness, richness and MRD, and MRD and temperature, and the negative β value, that the Ecological Limits hypothesis has influenced spatial patterns of shark diversity as defined by the framework constructed by H&S, at least at the global scale. This conclusion can be extended somewhat to the Lamniformes and Carcharhiniformes orders, though the strength of the influence in both clades is diminished compared to shark richness as a whole. We also conclude, based upon the lack of agreement between the regional analyses, that the ELH alone is insufficient to account for shark biodiversity patterns at smaller scales, and the patterns observed are more likely to be attributed to differences in geography. Correlations between richness and energy gradients alone are insufficient for differentiating between biogeographical and ecological hypotheses, and examining multiple spatial and phylogenetic metrics should better enable researchers to assess the comparative influence of different theories.

These results might be improved upon with the use of a mid-domain model in order to examine the results expected by randomly placed ranges. The simulation provided by Hurlbert and Stegen (2014b) could be made more complex in order to more adequately capture the specifics of the taxa in question, such as accounting for actual branch lengths in phylogenetic trees, examining nonlinear relationships between metrics, and including more spatial dimensions. It may also be useful to examine Chondrichthyans through the Metabolic Niche framework (McClain *et al*., 2012); partitioning shark species between high energy and low energy taxa and subsequently examining how each group interacts with the energy proxy may lend further support to the idea that the global distribution of species is determined in part by how much energy is available to a region and how precisely organisms use that energy with regards to their individual niche.

## Acknowledgments

College of Charleston School of Science and Math provided funding to ERS and DJM. The iDiv Biodiversity Synthesis lab group and the Naylor lab provided helpful feedback on the development of this project.

## Biosketch

Emmaline Sheahan performs research on drivers of diversity over large spatial and temporal scales in order to understand the general rules governing how species accumulate and change. She is primarily interested in the ecology and evolution of Chondrichthyes.

## Data Availability Statement

The spatial polygon range data used in this research is publicly available on Zenodo (https://doi.org/10.5281/zenodo.6321610).

