## Supplemental Figures for "Global shark species richness is more constrained by energy than evolutionary history"

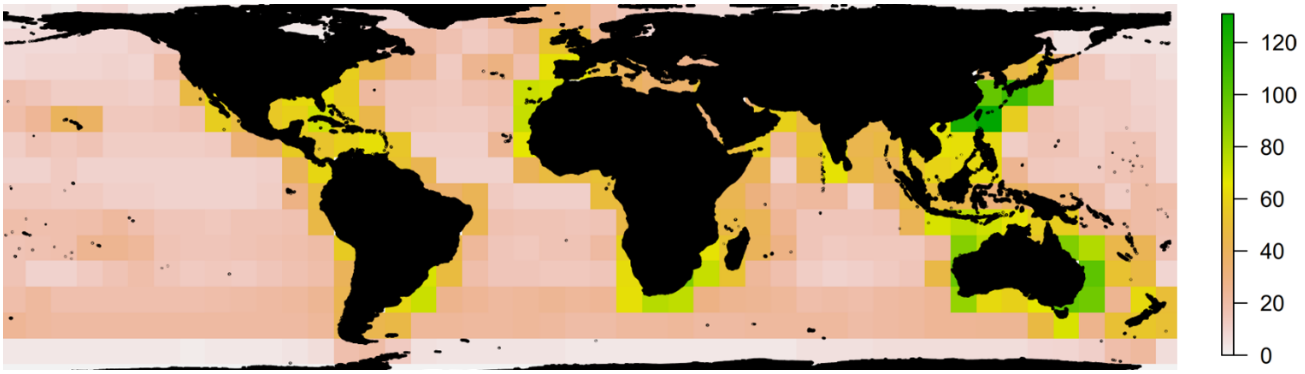


**Supplemental figure 1.** A map of global shark richness for all 534 species used in the analysis at an 880 x 880 km resolution. The greener the pixel, the higher the number of species in the pixel. Richness generally is more concentrated on the coasts, with hotspots around South Africa, Eastern and Western Australia, and Southern Japan.

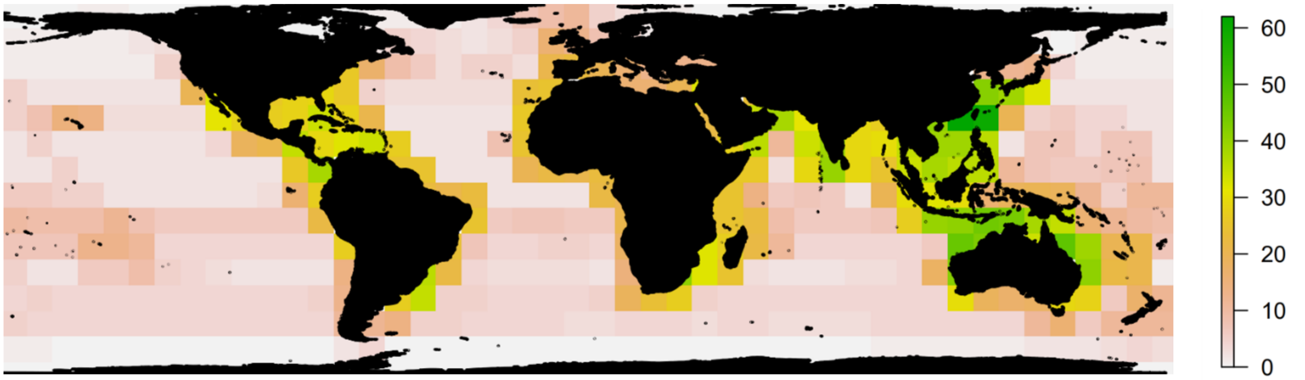


**Supplemental figure 2.** A map of ground shark richness for the 272 species used in the subclade analysis at the 880 x 880 km scale. The greener the pixel, the higher the number of species in the pixel. Ground shark richness is largely coastal and appears to be more concentrated in the Eastern hemisphere.

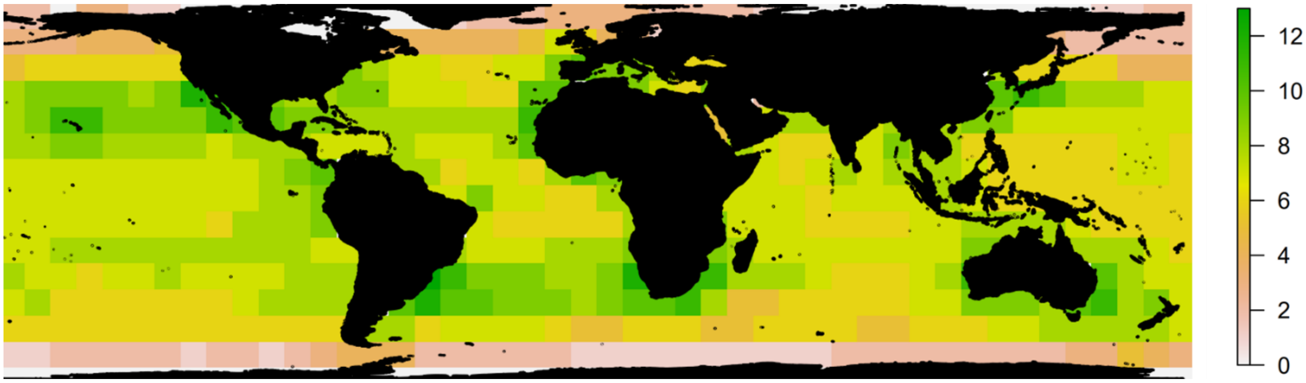


**Supplemental figure 3.** A map of mackerel shark species richness for the 15 species used in the subclade analysis. The greener the pixel, the more species there are in the pixel. Mackerel sharks exhibit more of a pelagic signal than the ground sharks and global analysis.

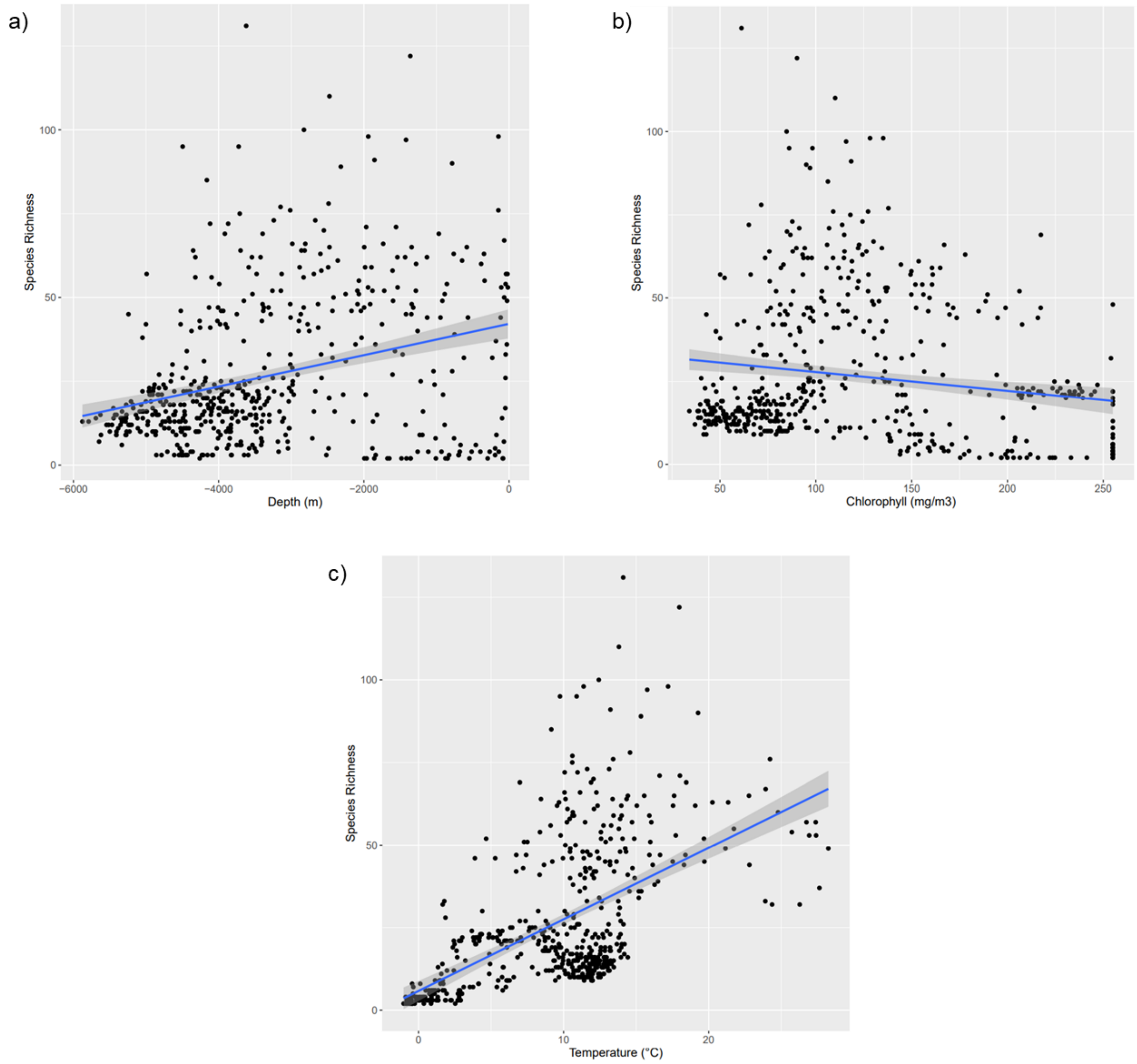


**Supplemental figure 4.** Scatter plots of a) species richness versus depth, b) species richness versus chlorophyll, and c) species richness versus temperature taken from rasters at the 880 x 880 km resolution. Because shark species richness is most strongly correlated with temperature, we used that environmental variable as our energy proxy.

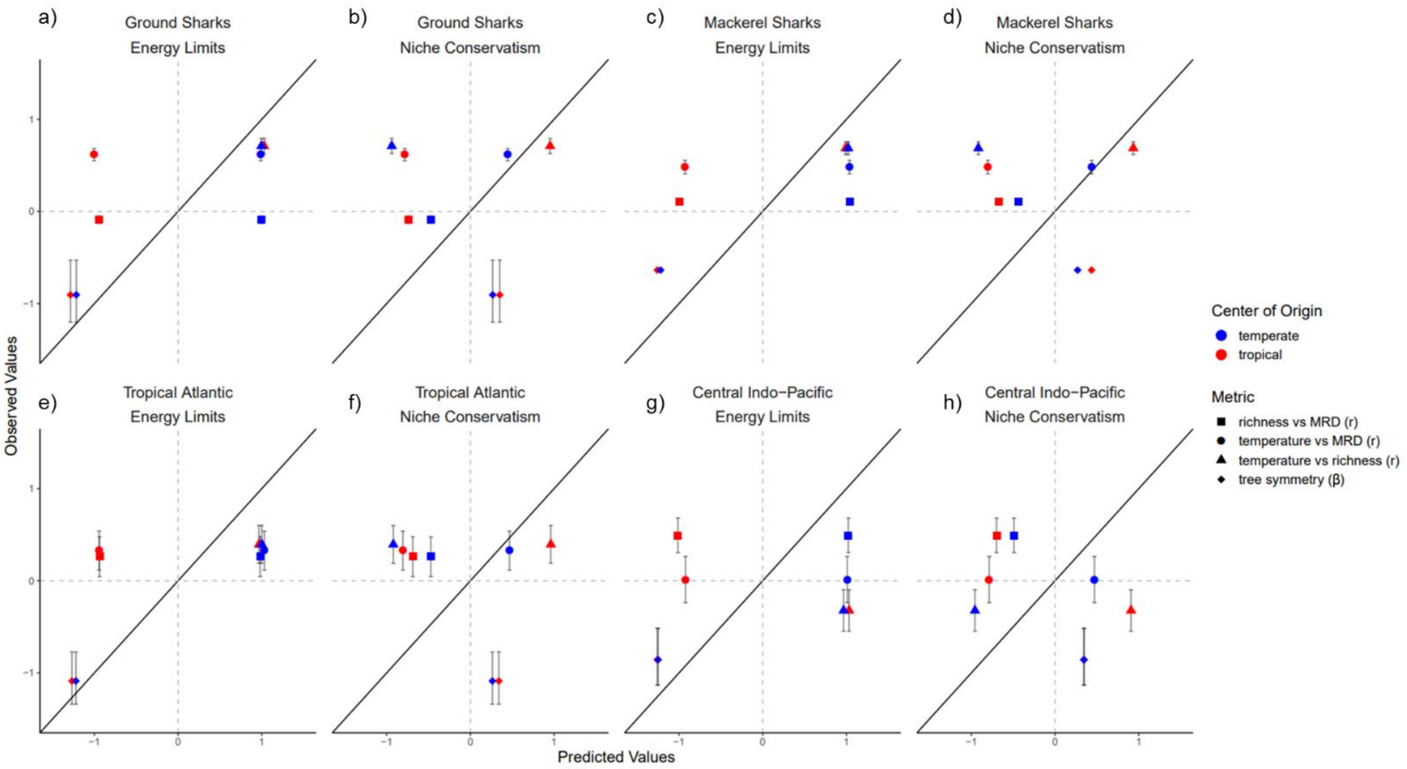


**Supplemental figure 5.** Predicted vs observed plots for the global analysis of the four metrics for a) ground sharks under ecological limits, b) ground sharks under niche conservatism, c) mackerel sharks under energy limits, d) mackerel sharks under niche conservatism, e) the Tropical Atlantic realm under energy limits, f) the Tropical Atlantic realm under niche conservatism, g) the Central Indo-Pacific realm under energy limits, and h) the Central Indo-Pacific realm under niche conservatism. The solid line is the one-to-one line; the closer the points are to the line, the better the fit between the model and reality. The 95% confidence interval of the observed metrics is provided for each metric, but it is sometimes obfuscated by the plotting symbol. The predicted values have been randomly jittered slightly to decrease point overlay. Points representing a tropical center of origin are red, and points representing a temperate center of origin are blue.
